# Mutational processes of tobacco smoking and APOBEC activity generate protein-truncating mutations in cancer genomes

**DOI:** 10.1101/2023.03.19.533271

**Authors:** Nina Adler, Alexander T. Bahcheli, Kevin Cheng, Khalid N. Al-Zahrani, Mykhaylo Slobodyanyuk, Diogo Pellegrina, Daniel Schramek, Jüri Reimand

## Abstract

Mutational signatures represent a footprint of tumor evolution and its endogenous and exogenous mutational processes. However, their functional impact on the proteome remains incompletely understood. We analysed the protein-coding impact of single base substitution signatures in 12,341 cancer genomes from 18 cancer types. Stop-gain mutations (SGMs) were strongly enriched in the signatures of tobacco smoking, APOBEC cytidine deaminases, and reactive oxygen species. These mutational processes affect specific trinucleotide contexts to substitute serine and glutamic acid residues with stop codons. SGMs are enriched in cancer hallmark pathways and tumor suppressors such as *TP53, FAT1*, and *APC*. Tobacco-driven SGMs in lung cancer correlate with lifetime smoking history and highlight a preventable determinant of these harmful mutations. Our study exposes SGM expansion as a genetic mechanism by which endogenous and carcinogenic mutational processes contribute to protein loss-of-function, oncogenesis, and tumor heterogeneity, providing potential translational and mechanistic insights.

## INTRODUCTION

Cancer is driven by a few somatic mutations that enable oncogenic properties of cells, however most mutations in cancer genomes are functionally neutral passengers ^1,2^. Somatic mutations are caused by endogenous and exogeneous mutational processes with complex context- and sequence-specific activities that collectively mark tumor evolution and exposures over time ^3^. Single base substitution (SBS) signatures are the indicators of mutational processes in cancer genomes that can be inferred through a computational decomposition of somatic single-nucleotide variants (SNVs) and their trinucleotide sequence context in large cancer genomics datasets ^4,5^. SBS signatures have been linked to clock-like mutational processes of aging ^6^, deficiencies in DNA repair pathways ^7^, endogenous mutational processes such as the activity of APOBEC cytidine deaminases ^8^, environmental carcinogens such as UV light ^9^, lifestyle exposures such as tobacco smoking ^10^, dietary components such as aristolochic acid ^11^, as well as the effects of cancer therapies ^12,13^. The causes of other signatures remain uncharacterised. Mutational signatures are also increasingly found in the somatic mutation profiles of healthy tissues, indicating that the mutational processes contribute to mutagenesis in normal and pre-cancerous cells ^14,15^. Individual driver mutations in cancer genomes have been attributed to the activity of certain mutational processes ^16,17^. While some mutational signatures identified in cancer genomes can be reproduced in experimental systems ^9,18,19^, their mechanistic and etiological characterization is an ongoing challenge. As mutational processes are thought to predominantly generate passenger mutations, their broad functional implications on protein function and cellular pathways remain incompletely understood.

Here we hypothesized that the mutational processes of SNVs have specific impacts on protein-coding sequence due to their trinucleotide sequence preferences encoded in SBS signatures. By characterizing the co-occurrence of mutational signatures and the sequence impact of associated SNVs in thousands of cancer genomes, we find that nonsense SNVs corresponding to stop-gain mutations (SGMs) are significantly associated with specific mutational processes of tobacco, APOBEC, and reactive oxygen species. SGMs are the most impactful class of SNVs that cause premature stop codons and result in truncated proteins or nonsense-mediated decay. Some consequences of these mutational processes appear as driver mutations in tumor suppressor genes and hallmark cancer pathways. These processes represent preventable carcinogenic exposures as well as endogenous sources of DNA damage, and their activity is explained by their sequence-specific interactions with the genetic code of stop codons. Our report provides direct evidence of the functional genetic impact of mutational signatures in cancer genomes and their interactions with the molecular and lifestyle drivers of the mutational processes, suggesting a role for these signatures in tumor heterogeneity and progression.

## RESULTS

### Protein-truncating mutations in cancer genomes are enriched in mutational signatures of tobacco smoking, APOBEC, and ROS

To study the protein-coding impact of SBS signatures, we analysed 12,341 cancer genomes from 18 major tissue sites using data in three pan-cancer cohorts: The Cancer Genome Atlas (TCGA) PanCanAtlas ^20^ with 6509 exomes, Pan-Cancer Analysis of Whole Genomes ^4^ (PCAWG) with 2360 whole genomes, and Hartwig Medical Foundation ^21^ (HMF) with 3472 whole genomes (**Figure 1a**) (**Supplementary Figure 1**). Hypermutated and low-confidence samples were filtered. 1.75 million exonic SNVs were classified based on their protein-coding function as missense (67.4%), silent (27.7%), stop-gain (4.6%), stop-loss (0.1%) and start-loss (0.1%) mutations. We used consensus mutational signature calls of PCAWG ^4^ and annotated the signatures in the TCGA and HMF datasets using the SigProfiler software ^5^. Using these three datasets allowed us to replicate our findings across sequencing platforms, variant calling pipelines, and signature analysis methods. We performed a mutation enrichment analysis by asking which specific mutational signatures were found in the five functional SNV classes significantly more often than expected from chance alone. Systematic analysis of the 18 cancer types in the three genomics datasets revealed 332 associations of mutational signatures and protein-coding variant function (Fisher’s exact test, FDR < 0.01) (**Supplementary Figure 2**).

**Figure 1.**
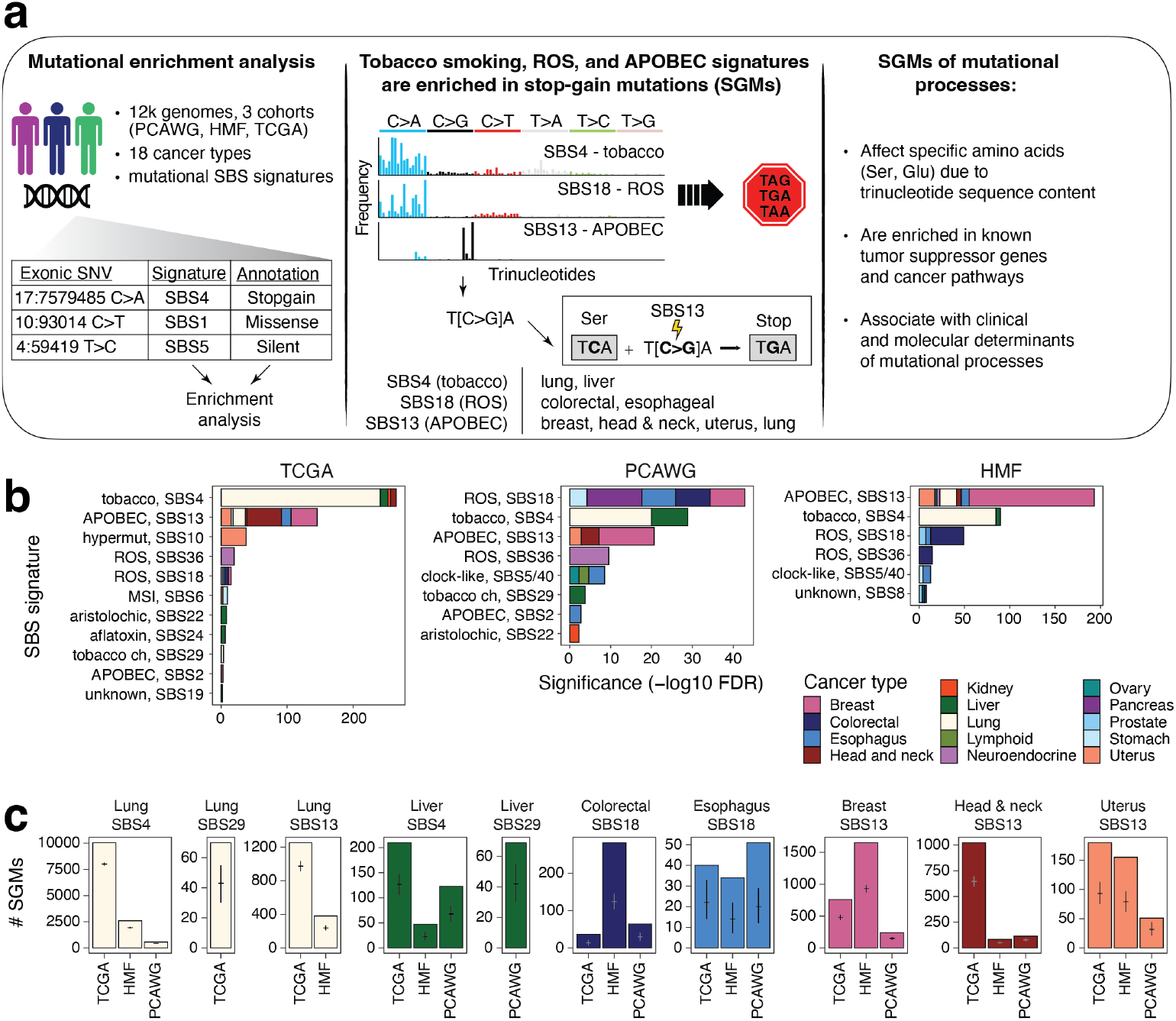
Protein-coding impact of mutational signatures in cancer genomes and associations with stop-gain mutations (SGMs). **(a)** Overview of study. Left: The associations of protein-coding impact of somatic single-nucleotide variants (SNVs) and the mutational signatures of single base substitutions (SBS) were studied using enrichment analysis in >12,000 cancer genomes. Middle: SGMs were enriched in the SBS signatures of tobacco smoking, APOBEC and ROS. The enrichments are explained by the trinucleotide preferences of the mutational processes that affect the genetic code of certain amino acids, converting these to stop codons. Right: Mutational signatures of SGMs were further studied in the context of affected driver genes and pathways as well as the clinical and molecular correlates of the mutational processes. **(b)** Significant enrichments of mutational signatures in SGMs in multiple cancer types and in three genomics datasets (FDR < 0.01). Bar plots show the cumulative significance of enriched SBS signatures in SGMs in various types of cancer. Tobacco smoking, APOBEC activity, and ROS exposure are the major mutational processes that contribute SGMs. **(c)** Observed and expected counts of SGMs derived from the most significant mutational processes in the three datasets of cancer genomes (TCGA, HMF, PCAWG). Mean expected mutation counts with 95% confidence intervals (CI) from binomial sampling are shown on the bars.

We focused on stop-gain mutations (SGMs) (i.e., nonsense SNVs), the most disruptive class of SNVs that induces protein truncations and loss of function (LoF). SGMs were consistently enriched in the SBS signatures of three major mutational processes of tobacco smoking, APOBEC activity, and reactive oxygen species (**Figure 1b-c**). First, the tobacco smoking signature SBS4 with frequent C>A transversions ^22^ was enriched in SGMs in primary lung cancers in TCGA (10,054 *vs*. 8,006 expected SGMs, fold-change (FC) = 1.26, FDR = 4.6 x 10^-242^; Fisher’s exact test) and metastatic lung cancers in HMF (FC = 1.34; FDR = 1.9 x 10^-85^). Similarly, SGMs were also enriched in the SBS4 signature in the three cohorts of liver cancer samples (FDR < 10^-5^). The SBS29 signature attributed to tobacco chewing was also associated with SGMs in lung and liver cancers (FDR < 0.001).

Second, the APOBEC signature SBS13 was enriched in SGMs in multiple cancer types, especially in breast (1653 SGMs observed *vs*. 931 expected, FDR = 1.1 x 10^-138^, HMF), head & neck (FC = 1.58; FDR = 3.3 x 10^-53^; TCGA), uterine, lung, and esophageal cancers. Notably, SBS13 appeared as the predominant APOBEC signature of SGMs while the alternative APOBEC signature SBS2 was not enriched in SGMs. SBS2 and SBS13 both preferably affect TCN trinucleotides, however SBS13 is primarily characterised by C>G and C>A mutations, while C>T mutations are common to SBS2, explaining the preferential enrichment of SBS13 to convert TCN to stop codons (TAG, TAA, TGA).

Third, SBS18 and SBS36, the two mutational signatures associated with reactive oxygen species (ROS), were also enriched in SGMs. These SBS signatures characterised by C>A mutations were especially enriched in cancers of the digestive system such as colorectal cancer (HMF: 282 SGMs observed *vs*. 124 expected; FDR = 5.0 x 10^-37^), esophageal cancer and stomach cancer, as well as pancreatic, neuro-endocrine and breast cancers. As ROS signatures were overall less frequent in cancer genomes than other signatures, fewer SBS18-associated SGMs were also found. Less-frequent carcinogenic signatures of aflatoxin and aristolochic acid exposures were also significantly associated with SGMs. These observations were consistent in primary and metastatic cancers, and their detection in independent WGS and WES datasets also lends confidence to our findings.

We validated the associations of SGMs and SBS signatures with additional analyses. First, we repeated the enrichment analysis using a probabilistic approach that accounted for all potential sample-specific signature exposures for annotating individual SNVs. By sampling these signature annotations repeatedly, we confirmed that SGMs remained highly significantly enriched in the SBS signatures of tobacco smoking, APOBEC, and ROS (**Supplementary Figure 3**). The probabilistic analysis also showed an even stronger enrichment of APOBEC signatures in SGMs in lung cancer compared to the analysis of top signature annotations, while the highly significant enrichment of SGMs in the smoking signature was somewhat attenuated in the probabilistic analysis. Previous studies indicate that both the tobacco smoking and APOBEC processes contribute somatic mutations in lung cancer whereas APOBEC is more involved in later mutagenesis ^23^, potentially explaining this observation. Accordingly, a subset of SGMs were likely attributable to either tobacco smoking or APOBEC signatures, or the age-associated signature SBS5 (**Supplementary Figure 4**). Second, we performed a pan-cancer analysis by combining samples of all cancer types and again recovered the tobacco smoking, APOBEC and ROS signatures with very strong enrichments of SGMs (**Supplementary Figure 5**). Thus, the exogenous mutational process of tobacco smoking and the endogenous processes of APOBEC activity and ROS appear as major drivers of disruptive protein-truncating mutations that may directly affect protein function and regulation in cancer.

We also reviewed the enriched SBS signatures of missense and silent SNVs (**Supplementary Figure 2**). Silent mutations were enriched in the mitotic clock-like signature SBS1 consistently in most cancer types across the three cohorts. SBS1 includes C>T transitions caused by 5-methylcytosine deamination. In contrast, missense SNVs were often enriched in the common clock-like signatures SBS5 and SBS40 that have relatively featureless (flat) trinucleotide profiles. Associations with APOBEC signatures were also identified in multiple cancer types: SBS2 was often enriched in silent SNVs while SBS13 was enriched in missense SNVs. Silent and missense SNVs comprise a large variety of trinucleotides and codons, and these signatures are detected in many cancer types, potentially explaining these broad associations and suggesting that functional subclasses of missense mutations should be considered in future analyses. Also, sample sizes and signature exposure determine the statistical power for detecting associations of SBS signatures and SNV annotations in the various cohorts. Taken together, these results exemplify the complex landscape of functional impacts mutational processes enact on the protein-coding genome.

### SGM signatures and the genetic code

To explore the genetic mechanism underlying the enrichments of SGMs in the three major mutational processes, we studied the types of amino acids most frequently substituted by stop codons, focusing on tobacco smoking, APOBEC, and ROS signatures in lung, breast and colorectal cancers. Several types of amino acids were surprisingly frequently replaced by stop codons. Glutamic acid (Glu) residues showed the strongest enrichment of stop codon substitutions in all three SBS signatures (**Figure 2a**). In lung cancer, Glu>Stop substitutions were enriched four-fold in the tobacco smoking signature SBS4 compared to other signatures (43.8% vs 10.6%, FDR < 10^-300^). Glutamic acid residues were also significantly affected by the APOBEC signature in breast cancer and ROS signatures in colorectal cancer (FC > 3.6, FDR < 1.7 x 10^-57^). As expected, these enrichments are also supported by reference SBS signatures of the COSMIC database ^24^ (**Figure 2b, Supplementary Figure 6**). Based on cosine similarity scores (COS), the SNV trinucleotide profiles corresponding to Glu>Stop substitutions in our data were considerably more similar to the COSMIC reference SBS signature profiles of tobacco smoking SBS4 and ROS SBS18 (lung: COS_SBS4_ = 0.40; colorectal: COS_SBS18_ = 0.62), compared to the frequent clock-like signatures SBS5 and SBS40 that we used as controls (lung: COS_SBS5_ = 0.13; colorectal: COS_SBS40_ = 0.33). Besides Glu>Stop substitutions, other amino acids were also enriched in stop codon substitutions in tobacco smoking and ROS signatures, including glycine (Gly; 13.7%) and cysteine (Cys; 6.4%) residues (all FC>4.8; FDR < 3.6 x 10^-60^), while arginine, glutamine and other residues were less frequently affected by SGMs than expected.

**Figure 2.**
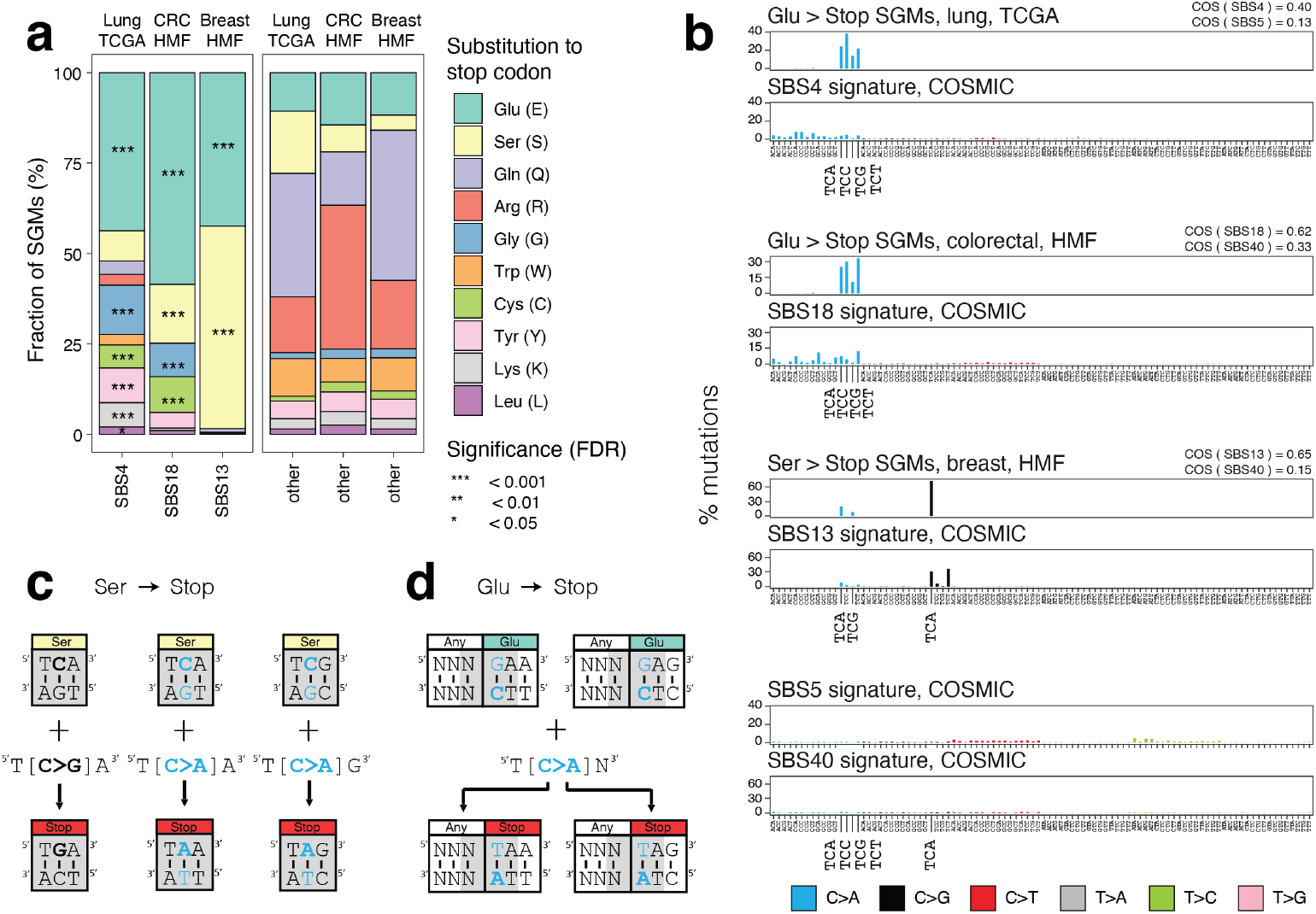
SBS signatures induce protein-truncating mutations by targeting the genetic code. **(a)** SGMs of tobacco smoking, APOBEC, and ROS signatures are enriched in specific amino acids. Stacked bar plots show the proportion of SGMs involved in various amino acid substitutions. Left: SGMs of the three major signatures: SBS4 of tobacco smoking in lung cancer, SBS18 of ROS in colorectal cancer, and SBS13 of APOBEC activity in breast cancer. Right: control SGMs associated with all other mutational signatures in these cancer types. Enrichment of signature-associated SGMs relative to controls are shown as asterisks (Fisher’s exact tests). **(b)** Trinucleotide profiles of SGMs reflecting substitutions of glutamic acid and serine residues in proteins (Glu > Stop, Ser > Stop) and the reference COSMIC signatures for tobacco smoking (SBS4), ROS (SBS18), and APOBEC (SBS13). As controls, the profiles of the most frequent mutational signatures are shown (SBS5/40; two plots at the bottom). Cosine similarity (COS) scores are used to compare the signature-associated SGMs and the COSMIC reference signatures (top right of each facet). The trinucleotide substitutions relevant to panels (c) and (d) are highlighted on the X-axis. **(cd)** Interactions of the mutational signatures and the genetic code of stop codons. The trinucleotides encoding serine and glutamic acid residues are shown as rectangles. The trinucleotides that can create stop codons when mutated are shown in grey. The trinucleotide changes required to create these stop codons are characteristic of the mutational processes of tobacco smoking, APOBEC, and ROS. **(c)** Serine substitutions to stop codons. C>G and C>A transversions in SBS13 and SBS18 induce stop codons by substituting the middle nucleotides in the three codons of serines (yellow). **(d)** Glutamic acid substitutions to stop codons. C>A transversions in SBS4, SBS13, and SBS18 induce stop codons by affecting two consecutive codons (grey). Since SBS signatures are represented with pyrimidines as the reference nucleotides, this schematic shows the trinucleotides that are mutated to create stop codons from glutamic acids as their reverse complement sequences. Here, the reverse-complementary trinucleotide transversions substitute the two first nucleotides in the glutamic acid codon (teal) and the third nucleotide in any preceding codon (white). The trinucleotide changes shown are characteristic of the tobacco smoking signature SBS4.

The APOBEC signature SGMs in breast cancer encoded stop codon substitutions almost exclusively in serine and glutamic acid residues (55.9% and 42.4%, respectively). Ser>Stop substitutions were significantly more frequent in SBS13 compared to other signatures (4.2% expected; FC = 13, FDR < 10^-300^). Accordingly, the SNV trinucleotide profiles encoding Ser>Stop substitutions were considerably more similar to the COSMIC reference APOBEC signature (COS_SBS13_ = 0.65) than the more common SBS40 reference signature we used as a control (COS_SBS40_ = 0.15), confirming the mutational signature annotations of SGMs in our data. Lastly, ROS signatures were also enriched in Ser>Stop substitutions in colorectal cancer (FDR = 6.2 x 10^-6^) while no enrichment of SGMs in serine residues was seen in the tobacco smoking signatures in lung cancer.

To consolidate these statistical associations into a mechanistic model, we examined the genetic code of the most common stop codon substitutions involving serine and glutamic acid residues (**Figure 2b-c**). First, the APOBEC-associated Ser>Stop substitutions in breast cancer were predominantly encoded by T[C>G]A transversions (76.8%) as well as T[C>A]A and T[C>A]G transversions. The TCA and TCG trinucleotides encode the two serine codons that require one SNV to become stop codons. The corresponding stop codons TGA, TAA and TAG are induced by three of the transversions distinctive of the SBS13 APOBEC signature. Second, the Glu>Stop substitutions apparent in the tobacco smoking and ROS signatures were predominantly caused by T[C>A]N transversions that overlap two adjacent codons (**Figure 2d**). Here, the reverse-complementary trinucleotides NGA include the two first nucleotides of glutamic acid codons GAA and GAG, which are replaced with the stop codons TAA and TAG upon N[G>T]A transversions, respectively. This model explains the genesis of SGMs by the mutational processes of tobacco smoking, APOBEC and ROS.

### Driver genes and pathways with truncating mutations

To study the functional consequences of SGM signatures, we asked whether the mutations converge on specific genes. We focused on the six types of cancer in which consistent signature-SGM enrichments were found in the TCGA, PCAWG and HMF datasets. We identified 56 genes with significantly enriched SGMs of tobacco smoking, APOBEC, and ROS signatures compared to the exome-wide distributions of these signatures and all SGMs: 14 genes in lung and liver cancers with the tobacco smoking signature; 44 genes in breast, head & neck, and uterine cancers with the APOBEC signature; and one gene (*APC*) in colorectal cancers with the ROS signature (**Figure 3a-b; Supplementary Table 1**) (Brown FDR < 0.05, Fisher’s exact tests). The genes included 556 signature-associated SGMs in 467 cancer genomes in the combined datasets, representing 3.8% of all tumors we studied. A large fraction of these (24 genes or 43%) are known cancer genes of the COSMIC Cancer Gene Census (CGC) database ^25^, significantly more than expected from chance alone (24 found, 2/56 genes expected; P = 1.4 x 10^-20^). Several core tumor suppressor genes (TSGs) such as *TP53, FAT1, CDH1, RB1, NF1*, and *APC* were identified.

**Figure 3.**
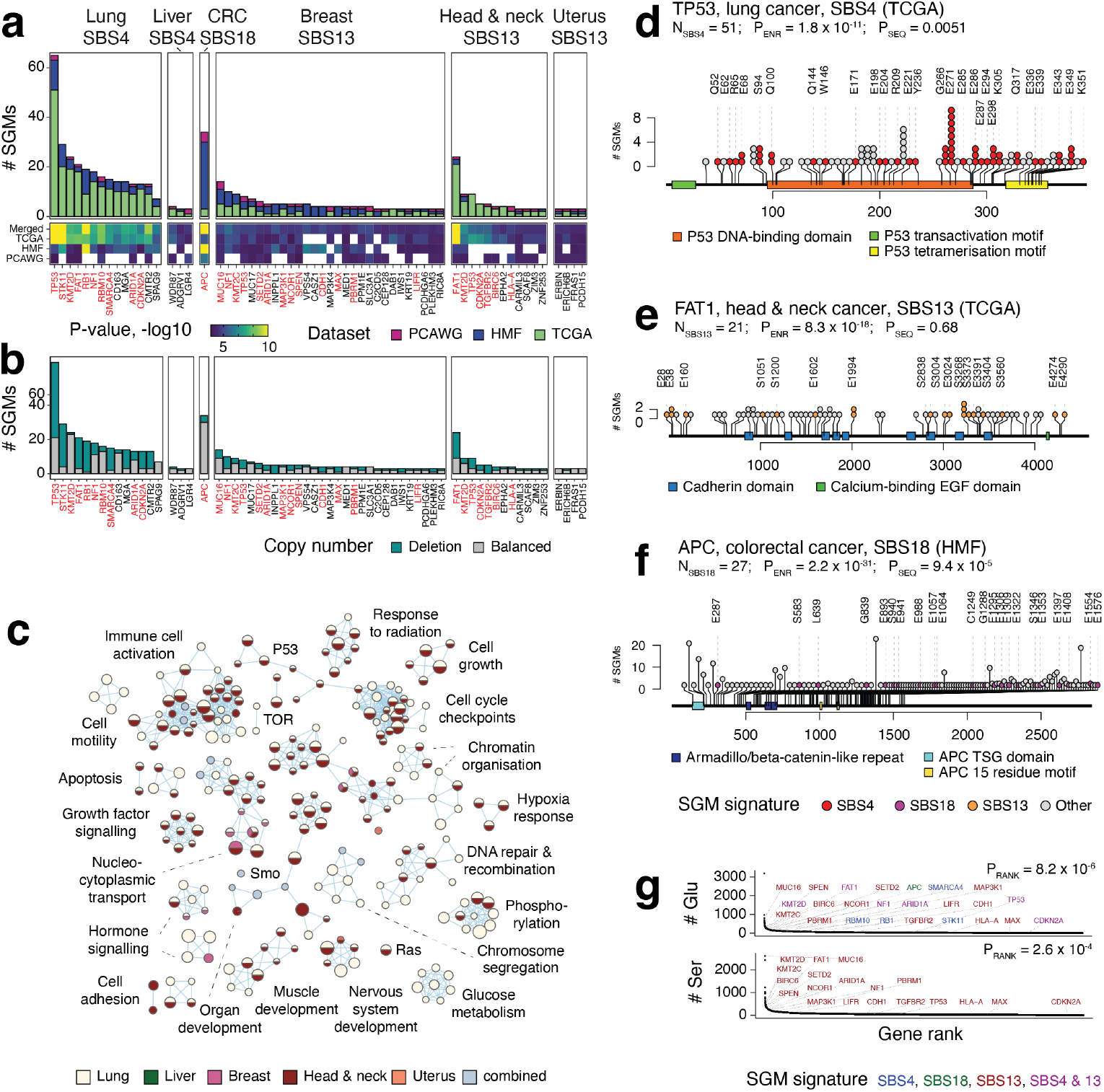
SGMs of tobacco smoking, APOBEC, and ROS signatures are enriched in tumor suppressor genes (TSGs) and cancer hallmark pathways. **(a)** Genes that are significantly enriched in SGMs driven by the three mutational signatures SBS4, SBS13 and SBS18. Each cancer type and cohort was analysed separately and the findings were integrated across the three cancer cohorts (Fisher’s exact tests, Brown FDR < 0.05). Known cancer genes are shown in red. **(b)** SGMs in the significantly enriched genes often co-occur with genomic copy-number losses in the same cancer samples, indicative of loss of heterozygosity. **(c)** Biological processes and molecular pathways with enriched SGMs of tobacco smoking and APOBEC signatures. Significant pathways were identified by merging evidence through the SGM signatures and the cancer types (ActivePathways, FDR < 0.05). The enrichment map is a network where enriched pathways and processes are shown as nodes and the edges connect the pathways that share genes. Nodes are colored by the cancer types in which the SGM enrichment was detected. Light blue represents the pathways that reached significance only when the evidence from the five cancer types was combined. **(d-f)** Examples of major TSGs that are enriched in SGMs driven by the mutational processes of tobacco smoking, APOBEC, and ROS. Colored circles show the signature-associated SGMs, their reference amino acid residues, and their sequence positions. PFAM protein domains are shown as colored rectangles. The number of SGMs of the mutational signature (N_SBS_), the enrichment P-value of SGMs (P_ENR_, Fisher’s exact test), and the P-value of SGMs accumulating towards either protein terminus (P_SEQ_, one-sample Wilcoxon rank-sum test) are shown. **(d)** SGMs in *TP53* in lung cancer are enriched in the tobacco smoking signature SBS4. **(e)** SGMs in *FAT1* in head & neck cancer are enriched in the APOBEC signature SBS13**. (f)** SGMs in *APC* are enriched in the ROS signature SBS18. **(g)** Genes enriched with SGMs of tobacco smoking, APOBEC, and ROS processes have a higher amino acid sequence content of Ser and Glu residues compared to all protein-coding genes (P_RANK_, one-sample Wilcoxon rank-sum test). Colors indicate the mutational signatures enriched in SGMs.

To further interpret the mutations functionally, we prioritised the genes with SGMs across cancer types, excluding colorectal cancer for which only one gene was found. Pathway enrichment analysis of SGM-ranked genes highlighted biological processes and pathways of such as apoptosis, growth factor signalling, cell motility, cell proliferation and development (ActivePathways ^26^ *FDR* < 0.05; **Figure 3c**). Most detected pathways were identified in multiple cancer types; primarily through the tobacco signature in lung cancer and the APOBEC signature in head & neck cancer. Thus, the SGMs generated by these mutational processes converge to tumor suppressor genes and cancer pathways and therefore contribute to loss of protein function, oncogenesis, and tumor progression.

The mutational signatures of tobacco smoking, APOBEC, and ROS were enriched in truncating mutations in important cancer genes. First, tobacco-associated SGMs were enriched in 14 genes in lung cancer across the three datasets, including *TP53*, in which most SGMs in the TCGA cohort (51/95) were driven by SBS4 (15 SBS4 SGMs expected, FDR = 1.8 x 10^-11^) (**Figure 3d**). Truncations in *TP53* preferentially occurred towards the disordered C-terminal tail involved in protein tetramerization (P = 0.0051, one-sample Wilcoxon rank-sum test) in which protein truncations have been associated with loss-of-function phenotypes of TP53 ^27^ and where post-translational modification sites are often mutated in cancer ^28^. SGMs of SBS4 and SBS13 showed high levels of functional activity in saturation mutagenesis of TP53 ^27^, supporting their roles in cancer phenotypes (**Supplementary Figure 7**). As another example, the second-ranking gene *STK11* had 20 of 22 SGMs attributable to the tobacco smoking signature in the TCGA dataset (1 SBS4 SGM expected; FDR = 5.6 x 10^-17^) (**Supplementary Figure 8**). Inactivating mutations of the *bona fide* TSG *STK11* (i.e., LKB1; serine/threonine kinase 11) are common in lung cancer, modulate differentiation and metastasis *in vivo*, and have been observed more frequently patients with smoking history ^29–31^. Smoking-associated truncations in STK11 accumulated towards the N-terminus of the protein (P = 0.002), suggesting that the mutational process of tobacco smoking directly contributes to early truncations and loss of function of this TSG. Similar enrichments of SGMs were seen in other core TSGs such as *RB1, NF1, ARID1A*, and emerging TSGs such as *MGA*, a transcription factor of the MYC network that suppresses growth and invasion in cellular and mouse models of lung cancer ^32^.

Second, SGMs of the APOBEC signature SBS13 were enriched in 44 genes in breast, head & neck, and uterine cancer. The most significant gene, *FAT1*, was found in head & neck cancer in TCGA and included 21 APOBEC-associated SGMs (1 SBS13 SGM expected, P = 8.3 x 10^-18^) (**Figure 3e**). *FAT1* encodes a proto-cadherin and master regulator of the Hippo pathway that controls organ growth, cell polarisation, and cell-cell contacts. *FAT1* is one of the most frequently mutated TSGs in cancer whose loss of function enhances tumor invasiveness, metastasis, and drug resistance ^33,34^, suggesting a link between APOBEC-induced protein truncations and disease outcomes. Interestingly, *FAT1* was also found in in lung cancer where SGMs were enriched in the tobacco signature SBS4. Besides *FAT1*, SGMs of the APOBEC signature were also seen in other hallmark cancer genes such *CDH1, TP53, CDKN2A* and *TGFBR2*, and putative cancer genes such as the receptor tyrosine kinase *EPHA2* that regulates glutamine metabolism in cancer through the Hippo pathway ^35^.

Third, the ROS signature SBS18 was enriched in SGMs in one gene, *APC*, which was identified in metastatic colorectal cancers of the HMF cohort (27/346 SGMs *vs*. 1 expected; P = 2.2 x 10^-31^) (**Figure 3f**). APC inactivation is an early oncogenic event that disrupts beta-catenin degradation and activates WNT signalling ^36,37^, suggesting a link of *APC* loss with the oxidative stress in the tumor microenvironment, diet, or with the therapies of metastatic cancers ^38^.

Tumor suppressor genes are often inactivated in cancer through multiple mechanisms. To determine whether TSGs were inactivated in samples with signature-associated SGMs, we asked whether the 56 SGM-enriched genes were also altered by genomic copy number (CN) losses in the same tumor samples (**Figure 3b**). Indeed, the SGM-enriched genes appeared to carry both protein-truncating mutations and copy-number losses in most relevant cancer samples (58.9% or 275/467). This was also apparent in individual TSGs such as *TP53* (67.1% or 55/82 samples), *STK11* (86.2% or 25/29) and *FAT1* (73.3% or 33/45). Thus, some SGMs contributed by the mutational processes of tobacco and APOBEC may be involved in biallelic inactivation of TSGs where one gene copy is inactivated by SGMs while the other copy is deleted.

We asked if the SGM-enriched genes were in accordance with our model of SGM signatures and the genetic code (**Figure 2c-d**). As expected, the proteins encoded by the 56 SGM-enriched genes encoded for significantly more glutamic acid residues relative to the reference human proteome (Wilcoxon rank-sum *P* = 8.2 x 10^-6^), while the subset of these proteins with APOBEC-associated truncations also associated with a higher serine content (*P* < 2.6 x 10^-4^) (**Figure 3g**). The genetic model was also supported at the level of individual genes. For example, most TP53 truncations of the tobacco smoking signature affected glutamic acids (32/51), while the APOBEC-associated truncations in FAT1 affected either serines (10/21) or glutamic acids (11/21). Furthermore, the same subset of core TSGs was enriched in both tobacco smoking- and APOBEC-associated SGMs in different cancer types (*e.g., TP53, CDKN2A, NF1, FAT1* and *ARID1A*), as the SGMs introduced through the two mutational processes converge across different cancer types. Thus, certain TSGs may be more vulnerable to these mutational processes and the resulting SGMs due to their protein sequence content, indicating an interplay of the mutational processes and positive selection against tumor suppressive function.

### Clinical and molecular associations of SGM signatures

We asked how the mutational processes of SGMs were manifested in individual cancer genomes. The largest number of SGMs was associated with the tobacco smoking signature in lung cancers. In TCGA, lung cancer samples included 10.5 tobacco smoking-associated SGMs per genome on average, whereas 73% of cancers had at least one and 39% had at least ten such protein-truncating mutations. In primary TCGA breast cancers, the mutational process of APOBEC was associated with an average of 1.1 SGMs and affected a quarter of samples. In metastatic breast cancers of the HMF dataset, APOBEC-driven SGM burden was higher (mean 2.3 SGMs per sample; 32% of samples), consistent with longer or higher APOBEC levels in advanced cancers ^39^. ROS-associated SGMs, while less frequent in cancer genomes overall, were most pronounced in metastatic colorectal cancers in HMF, affecting 23% of samples with an average of 0.5 SGMs per genome (**Figure 4a**). Therefore, a large fraction of cancer genomes has some SGMs and potential loss-of-function alterations through these mutational processes.

**Figure 4.**
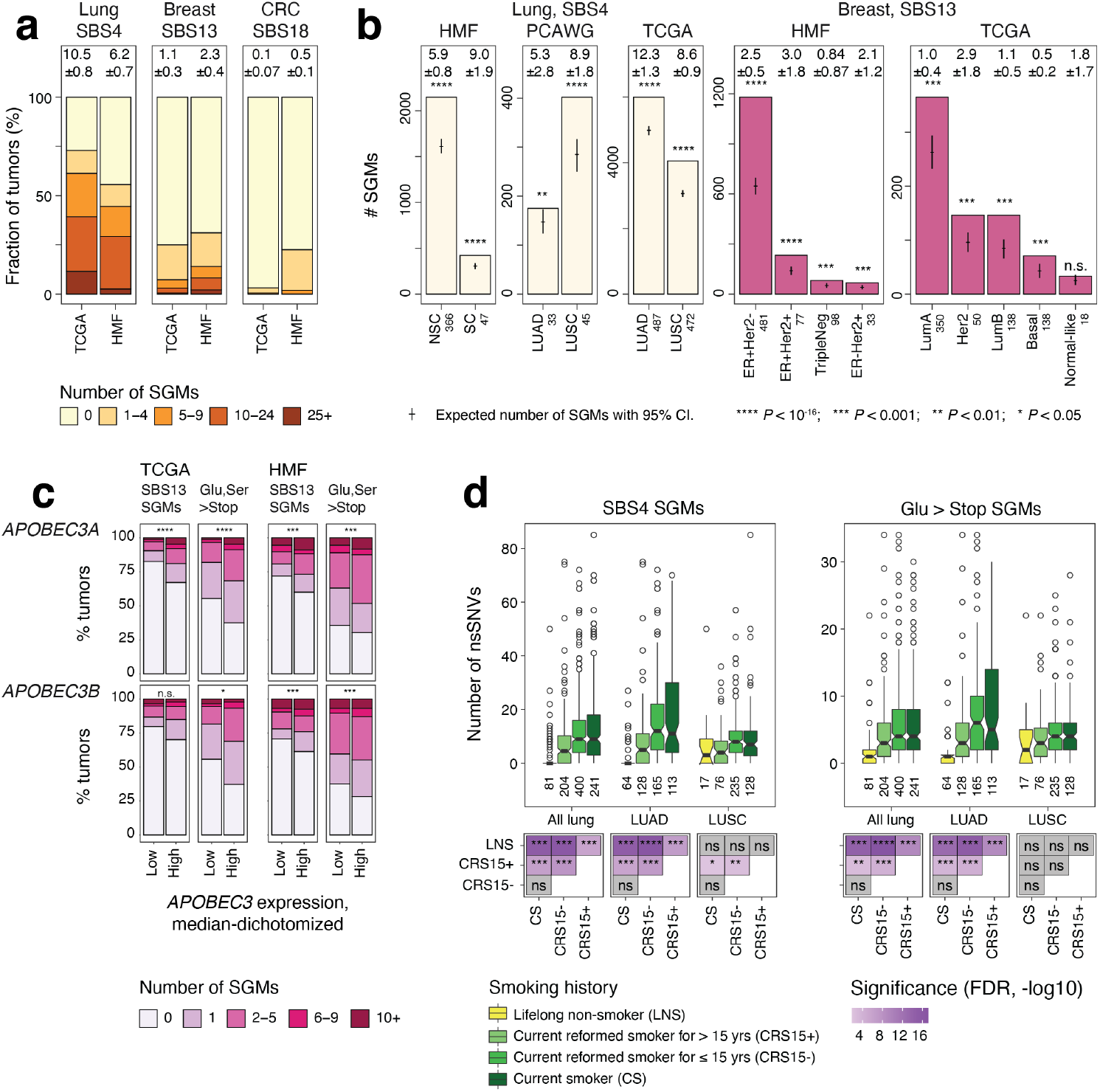
Molecular and clinical associations of SGMs with the mutational processes of tobacco smoking and APOBEC. **(a)** Mutational burden of SGMs of the three most significant mutational processes in cancer genomes. Stacked bar plots show the numbers of SGMs per cancer genome for the three SBS signatures: tobacco smoking in lung cancers (SBS4; left), APOBEC in breast cancer (SBS13; middle), and ROS in colorectal cancer (SBS18; right). Primary and metastatic cancer cohorts are compared (TCGA, HMF). Mean numbers of SGMs per cancer genome with ± 95% CI are shown above the bar plots. **(b)** SGMs of tobacco smoking and APOBEC signatures are also enriched in the molecular subtypes of lung and breast cancer. Expected total SGM counts with 95% CI are shown as points and whiskers. The counts of cancer samples in each group are shown in the X-axis labels. Lung cancer subtypes include small-cell (SC), non-small cell (NSC), adenocarcinoma (LUAD), and squamous cell carcinoma (LUSC) as annotated in the datasets. **(c)** Gene expression of *APOBEC3A* and *APOBEC3B* in breast cancer samples in TCGA associates with more frequent SGMs of the SBS13 signature (left) and more frequent Ser>Stop and Glu>Stop substitutions (right). The analysis compared cancer samples that were grouped by the expression of *APOBEC3A* and *APOBEC3B* genes (median-dichotomized; high *vs*. low). P-values of Poisson exact tests are shown. **(d)** The SGMs of the tobacco smoking signature are associated with the smoking history of lung cancer patients in TCGA. Box plots show the numbers of SGMs of the SBS4 signature (left) and Glu>Stop substitutions per cancer genome (right). The counts of cancer samples in each group are shown below the box plots. The statistical significance of correlating SGM burden with respect to smoking history is shown in the grid plot at the bottom (Wilcoxon rank-sum test, FDR-adjusted; * < 0.05; ** < 0.01; *** < 0.001; **** < 10^-16^).

We next studied the activity of SGM signatures at the level of cancer subtypes. The enrichment of SGMs in the tobacco smoking signature was detected in primary lung adenocarcinomas (LUAD) and squamous cell carcinomas (LUSC) in TCGA, as well as metastatic non-small cell and small cell cancers in the HMF dataset (**Figure 4b**). APOBEC associations with SGMs in breast cancer were also confirmed in the major histological and molecular subtypes of the disease and were also detected in primary breast cancers and metastatic cancers originating from the breast. Notably, the Her2-positive breast cancer subtype in TCGA had three-fold more APOBEC-driven SGMs than all other subtypes combined (2.9 vs. 0.95 SGMs per genome) (**Figure 4b**), consistent with earlier studies showing higher APOBEC activity in that subtype ^8^. Thus, the mutational processes generating SGMs are active across lung and breast cancer subtypes and in primary and metastatic cancers.

We analysed the mRNA abundance of APOBEC enzymes to study the molecular drivers of these mutational processes, focusing on breast cancer cohorts where most consistent signals of APOBEC-driven SGMs were found. In TCGA, the cancer samples with higher *APOBEC3A* or *APOBEC3B* expression had significantly more APOBEC-driven SGMs compared to cancers with lower expression (P = 2.1 x 10^-28^, Poisson exact test). Similarly, significantly more SGMs involving serine and glutamic acid residues were found (P = 2.0 x 10^-24^). The positive association of APOBEC enzyme expression and SGM burden was also observed in the HMF cohort of metastatic cancers (**Figure 4c**), as well as other APOBEC3 genes (**Supplementary Figure 9**) The association of APOBEC mutagenesis with gene expression of the two enzymes confirms earlier studies of cancer genomes and experimental systems ^19,40^ and connects the mutational processes of SGMs with the expected molecular pathway.

Lastly, we assessed SGMs in the context of smoking history of lung cancer patients in TCGA. Compared to patients with smoking history, the cancer genomes of lifelong non-smokers had fewer tobacco-associated SGMs of SBS4 (FDR < 10^-6^) and fewer Glu>Stop substitutions (FDR < 10^-9^). No significant differences in SBS4 SGM burden were found between current and recently-reformed smokers (FDR = 0.93); however, both groups had significantly more SBS4 SGMs than lifelong non-smokers and long-term reformed smokers (FDR < 10^-6^). Cancer subtype analysis confirmed the association between lifetime smoking and the burden of SBS4 SGMs in LUAD, while weaker signals were observed in LUSC (**Figure 4d**). This is expected as the LUAD subtype is more common among non-smokers than LUSC ^41^ (13.6% and 3.7% in TCGA, respectively) and the more varied composition of the LUAD cohort may contribute to the more pronounced association with SGMs. Therefore, SGMs in lung cancer genomes can be attributed to lifetime smoking activity, indicating a preventable cause of these impactful genetic aberrations. Overall, the clinical and molecular associations of mutational processes and SGMs provide insights to tumor heterogeneity and patient outcomes and have potential implications to biomarker and therapy development.

## DISCUSSION

Our pan-cancer analysis shows that the mutational processes of tobacco smoking, APOBEC, and ROS are a source of protein-truncating mutations in cancer genomes. The trinucleotide context of these mutational processes results in substitutions of glutamic acid and serine residues to stop codons, explaining the strong statistical associations observed in many cancer types. In support of this mechanism, we present several lines of evidence. First, the tumor suppressors with the strongest enrichments of SGMs also have a high protein sequence content of these amino acids. Second, we can identify the mutational processes of SGMs in large cohorts of primary and metastatic cancers of various disease types, and in whole-genome and whole-exome sequencing datasets. Third, the mutational burden of SGMs correlates with the molecular drivers of the mutational processes, including lifestyle tobacco exposure of lung cancer patients and the expression levels of APOBEC genes.

Our analysis ties together the functional impact of mutational processes and positive selection in cancer genomes. The genes with the most frequent SGMs associated with the three mutational processes are clearly enriched in core tumor suppressor genes, including early oncogenic drivers such as *APC*, later drivers of tumor progression and metastasis such as *CDH1*, as well as less-characterized cancer genes for further studies such as *EPHA2* and *MGA*. SGM-enriched genes converge to cancer hallmark pathways across multiple cancer types. In many cases, protein-truncations of TSGs are combined with copy-number deletions, indicating that SGMs contribute to biallelic inactivation. The trinucleotide preferences of the mutational processes imply that the TSGs with a higher protein content of glutamic acids and serines are more vulnerable to protein truncations caused by these mutational processes. In these genes, SGMs likely promote cancer development and are under positive selection.

The mutational processes that contribute SGMs are the major processes of somatic mutagenesis in many cancer types. Tobacco smoking appears as the most significant driver of SGMs in the cancers of lung, head and neck, and esophagus cancers that involve direct exposure to smoke. We also find SGM enrichments in lower-frequency carcinogenic processes of tobacco chewing, and the dietary carcinogens aflatoxin and aristolochic acids. Further, increased smoking is associated with higher SGM burden, indicating that the more an individual is exposed to tobacco smoke, the more likely they are to acquire protein-truncating somatic mutations in tobacco-exposed cells. Therefore, loss-of-function mutations in cancer genomes appear to be determined by lifestyle and environmental exposures of smoking and exposure to second hand-smoke.

APOBEC enzymes are core components of the innate immune system that are involved in restriction of virus replication and somatic antibody diversification ^42^. Aberrant APOBEC activity is a major cause of somatic mutations in many cancer types. APOBEC enzymes were described in virus restriction pathways that disable viruses by inducing missense and stop-gain mutations in viral RNA ^43^. This evolutionarily important mutational process of defense against pathogens corroborates our observations in cancer genomes. RNA editing by APOBEC1 causes a protein truncation in the apolipoprotein APOB that is required for lipid processing in the intestine ^44,45^. Somatic mutagenesis in normal human small intestines has been associated with APOBEC1 activity and includes protein truncations in TSGs ^46^. APOBEC mutagenesis is clustered in tissue-specific open-chromatin regions ^47,48^, indicating that the SGMs preferentially target actively expressed genes in which protein-truncating mutations are more likely to have functional consequences. Therapeutic APOBEC inhibition may help reduce the genesis of SGMs and the rein in tumor heterogeneity, especially as APOBEC has been linked to mutational processes, sub-clonal diversification, and driver mutations later in tumor evolution ^39^.

Oxidative stress and ROS are major sources of genomic instability that are associated with lifestyle factors common in developed countries, such as malnutrition, limited dietary antioxidant levels, obesity, and excess alcohol consumption ^49^. Oxidative stress is also associated with some anti-cancer therapies such as ionizing radiation and certain chemotherapeutic agents. SGMs of ROS signatures were especially apparent in metastatic colorectal cancers that are commonly treated with radiation therapy. Interestingly, rare cancer-predisposing germline mutations of the DNA repair enzyme MUTYH have been associated with the ROS signature SBS18 and more frequent SGMs ^50^, supporting our findings of SBS18-driven SGMs in cancer genomes. Thus, lifestyle variables, genetic makeup of patients and certain cancer treatments may contribute to loss-of-function mutagenesis, increasing genetic tumor heterogeneity and ultimately enabling additional molecular avenues of tumor progression and metastasis.

As cancers evolve, they become more heterogeneous and their paths to progression and metastasis become more varied. This heterogeneity is likely acquired through additional mutations that further deregulate cancer pathways and unlock therapy resistance. By inducing SGMs, the mutational processes of tobacco, APOBEC and ROS directly contribute to tumor heterogeneity by causing protein loss-of-function mutations. While not all these SGMs occur in core TSGs and directly drive cancer phenotypes, SGMs may involve genetic interactions with the core driver genes. Synergistic interactions may provide additional context-specific advantages to tumors in cases where the SGMs disrupt protective mechanisms and thereby enhance the phenotypes caused by core driver genes. On the other hand, SGMs may lead to synthetic lethal interactions where the SGMs disrupt a pathway that the core oncogenic pathway depends on. These interactions may be exploited for therapy by targeting other components of the SGM-disrupted pathway. Therefore, SGM-inducing mutational processes are likely to increase inter-and intra-tumoral heterogeneity through loss-of-function. Mutational processes of SGMs may also have implications on the tumor’s interactions with the immune system. SGMs lead to truncated and malformed proteins, some of which may be expressed on the cell surface and appear as neoantigens. Such truncated proteins may render the tumor more visible to the immune system, highlighting avenues for T cell-based immunotherapies. Deeper analyses of the proteogenomic impact of mutational processes, their etiology and genetic and lifestyle associations may lead to innovative biomarkers, mechanistic insights to cancer pathways, and leads for therapy development.

## METHODS

### Somatic mutations in cancer genomes

We analysed somatic SNVs in three cohorts of multiple cancer types: ICGC/TCGA Pan-cancer Analysis of Whole Genomes (PCAWG) ^2^ with whole genome sequencing (WGS) data of primary cancers, Hartwig Medical Foundation (HMF) ^21^ with WGS data of metastatic cancers, and the Cancer Genome Atlas (TCGA) PanCanAtlas ^20^ project with whole exome sequencing (WES) of mostly primary cancers. We used the Multi-Center Mutation Calling in Multiple Cancers (MC3) dataset of TCGA variant calls ^51^. We removed hypermutated tumor samples defined as those with >90,000 SNVs in WGS data and with >1800 SNVs in WES data, corresponding to genomes with approximately >30 SNVs/Mbps (n = 69 for PCAWG; n = 306 for HMF; n = 806 for TCGA). We excluded SNVs that did not pass the MC3 quality filter in TCGA. In WES data, we also removed lower-confidence samples with very few mutations for increased confidence in signature decomposition (<20 SNVs; 977 samples in TCGA). In HMF, we removed 140 duplicate cancer genomes of tumors of the same patients by selecting the sample with the highest tumor purity. We also removed 25 samples lacking HMF patient IDs. To enable analyses across the TCGA, PCAWG and HMF cohorts, cancer types were consolidated to 18 meta-types based on organs and/or anatomical sites, each with each cancer type including at least 25 samples in the three cohorts (**Supplementary Figure 1**). In HMF, the organ of the primary tumor was used for cancer type classification. Cancers of and less-frequent primary sites and of unknown origin (HMF) were excluded. In total, we analysed 1,751,110 exonic SNVs. The functional effects of SNVs on protein-coding genes were annotated using the ANNOVAR software ^52^ (version 2019-Oct-24)by using the canonical protein isoforms of the genes. The final dataset contained 12,341 cancer genomes (2360 in PCAWG, 3472 in HMF, 6509 in TCGA). This included some samples that were present in both the PCAWG and TCGA cohorts (n = 484). The duplicate samples in PCAWG and TCGA were retained to provide additional technical validation across the sequencing platforms, variant calling pipelines, and signature mapping strategies used to produce the datasets.

### SBS signatures

Mutational signatures for single base substitutions (SBS) in PCAWG were retrieved from the consensus PCAWG dataset ^4^. In HMF and TCGA datasets, we separately assigned known SBS signatures to SNVs using the SigProfilerSingleSample software (version 0.0.0.27) ^5^ and the COSMIC SBS signature catalogue (version 3) ^5,24^. For most analyses, each SNV was assigned to the most probable SBS signature based on these signature exposure prections. We removed a small subset of samples in WGS data that were potentially contaminated with sequencing artefacts as defined by the presence of more than 20% of SNVs assigned to SBS27, SBS43, and SBS45-SBS60, comprising nine samples in PCAWG and four samples in HMF. The TCGA dataset was not further filtered beyond the MC3 quality filter ^51^. To verify the tobacco, APOBEC and ROS signatures of SGMs in lung, breast and colorectal cancers respectively, we computed cosine similarity (COS) scores to evaluate the similarity of the SGM trinucleotide profiles with the reference SBS trinucleotide signatures of the COSMIC database. COS scores were separately computed for all SGMs and specific amino acid substitutions involving serines and glutamic acids. As controls, we also computed the equivalent COS scores comparing the SGM trinucleotide profiles with the two clock-like feature-less SBS signatures, SBS5 and SBS40, which were the most assigned SBS signatures in the respective cancer types.

### Enrichment analysis of protein-coding SNV classes and mutational signatures

We performed a comprehensive enrichment analysis of functional SNV annotations and mutational SBS signatures by separately comparing all consolidated cancer types in the three cancer cohorts. The analysis evaluated whether the classes of exonic SNVs (*i.e*., missense, stop-gain (i.e., non-sense), silent, start-loss, stop-loss) were significantly enriched in certain mutational signatures more often than expected from the independent binomial distributions of these SNV classes and the SBS signatures in all protein-coding regions of a given cancer type and cohort. For each cancer type, we tested the signatures that were reasonably frequently detected, had at least 100 SNVs per cancer type and cohort, and included at least one SNV of the tested variant annotation class (*e.g*., SGM), excluding signatures annotated as sequencing artefacts in the COSMIC database (see above). Certain signatures associated with the common mutational processes were combined: clock-like signatures SBS5 and SBS40 (SBS5/40), UV signatures SBS 7a/b/c/d (SBS7), hypermutation-associated signatures SBS10a/b (SBS10), and the signatures SBS17a/b (SBS17). Since this analysis focused only on protein-coding regions, we excluded SNVs outside exons in the WGS datasets from our statistical tests. To provide comparable analyses of WGS and WES datasets and reduce the inflation of significance in better-powered WGS datasets, we excluded non-exonic variants from the statistical tests. Statistical analysis was conducted using one-tailed Fisher’s exact tests that asked whether a set of SNVs derived from a given SBS signature and another set of SNV with a given functional annotation were overlapping significantly more often than expected by chance alone. The resulting P-values were adjusted for multiple testing using the Benjamini-Hochberg False Discovery rate (FDR) method ^53^. Results were considered significant if FDR < 0.01. Expected values of mutations sharing SBS signatures and functional annotations were sampled from the independent binomial distributions over 10,000 iterations, parametrized by the product of the probabilities of signature mutations and functional annotations, respectively. Using a similar approach, we also asked if specific types of amino acids were more likely to be substituted with stop codons through the SGMs driven by the identified SBS signatures. This analysis focused on only three cancer types and and three SBS signatures in the cohorts with the strongest signals (SBS4 in lung cancer in TCGA, SBS13 in breast cancer in HMF, SBS18 in colorectal cancer in HMF). Fisher’s exact tests were performed to assess whether certain amino acids were co-occurring with the signatures significantly more often than expected from the individual binomial distributions of the signature-associated variants and the substituted amino acid. The resulting P-values were corrected for multiple testing using FDR.

### Confirming the enrichment of SGMs in the major SBS signatures with probabilistic sampling

All major analyses in our study considered the most probable SBS signature for each SNV. To confirm our findings by accounting for the uncertainty in the signature annotations of individual SNVs and tumor samples, we performed a sampling analysis in which we assigned signatures to individual SNVs probabilistically over 100 iterations. Each SNV was assigned an SBS signature based on the multinomial distribution parametrised by the probabilities of all the SBS signatures identified in the given cancer genome. This procedure allowed the less-probable signatures to be included in the SNV annotation based on their probabilities. The 100 probabilistically sampled SNV-to-signature assignments were then systematically analysed using the enrichment analysis approach described above, to determine which signatures were enriched in SGMs in various cancer types. Significant results for each iteration were selected after iteration-specific multiple testing correction (FDR < 0.05). The fold-changes and FDR-values of the different iterations were then visualised as volcano plots that summarized fold change and FDR in all iterations.

### Analysis of SBS signatures and SGMs in genes

Genes with significant signature-associated SGMs were identified using one-tailed Fisher’s exact tests separately for the three major signatures (SBS4, SBS13, SBS18). The tests compared the distribution of SGMs of each SBS signature in a gene relative to the distributions of all SGMs and all mutations of that SBS signature in all protein-coding genes combined. This analysis only used exonic mutations and excluded mutations in non-coding regions, similarly to the exome-wide analysis described above. Fisher’s exact tests were conducted for each gene separately and in all three cohorts separately (TCGA, HMF, PCAWG). Genes were only tested if they had at least one SGM assigned to the given mutational signature. The resulting P-values for each gene were merged using the Brown procedure ^54^ and corrected for multiple testing using FDR. Significant genes were selected based on the Brown merged FDR-values (FDR < 0.05). Known cancer genes of the COSMIC Cancer Gene Census (CGC) database ^25^ (version 2020-09-17, accessed 2021-10-21) were highlighted in the resulting gene list. A Fisher’s exact test was used to determine whether the CGC genes were found in the list more often than expected, using all protein-coding genes as the background set. In an additional analysis, all protein-coding genes were ranked according to the numbers of glutamic acid (Glu) and serine (Ser) residues in their canonical protein isoforms. Genes identified in the SGM enrichment analysis from above were tested for higher-than-expected Glu and Ser content using one-tailed Mann-Whitney U-tests that determined whether the ranks of the selected genes were significantly higher than the median rank across the reference human proteome. For each candidate gene, we determined whether the sequence positions of the signature-associated SGMs were distributed towards either the N or C terminus of the protein more often than expected. One-tailed one-sample Wilcoxon rank-sum tests were used for this analysis. To analyse the functional impact of SGMs in TP53, we obtained data from saturation mutagenesis screens from the study by Giacomelli *et al*. ^27^ and compared the Z-scores of TP53 functional activity among four classes of SNVs: (i) SGMs associated with SBS4 and/or SBS13 in any cancer sample in our datasets (i.e., PCAWG, TCGA, HMF combined), (ii) all other SGMs of other SBS signatures (i.e., excluding SBS4 and SBS13) observed in any cancer sample in our datasets, (iii) missense SNVs observed in any cancer sample in our datasets, and (iv) as controls, all other mutations of TP53 studied in the mutagenesis screens but not in any human cancer genomes. Only unique mutations were analysed. Statistical significance estimates between the groups were determined using Wilcoxon rank-sum tests.

### Analysis of copy number alterations (CNAs) and SGMs

We aimed to identify potential biallelic inactivation cases where the gene was disrupted by both SGMs and copy number (CN) alterations leading to the genomic losses of the gene in the same tumor. We studied the 56 genes with significantly enriched signature-associated SGMs from our analysis that included 556 SGMs in 467 tumors in total. Separate strategies to select CNAs were used for the TCGA dataset and the PCAWG and HMF datasets. For TCGA samples profiled previously using SNP6 microarrays, we analysed the relative digital somatic CN calls of each gene as from previous consensus datasets. Gene losses in TCGA were defined through gene CN < 0. For PCAWG and HMF samples previously profiled using WGS, we analysed the CN values of genomic segments defined in these projects. To define the CN value for each gene, we considered the overlapping genomic segment with the lowest CN and of at least 1 kbps in length. To define gene losses in PCAWG and HMF, we used different criteria for autosomes and the X chromosome, and for samples with and without potential whole-genome duplication (WGD) events. A cancer genome was predicted to have undergone WGD if the genome-wide CN > 2.5. For non-WGD samples, we defined gene losses in autosomes through gene CN < 1.5. For WGD samples, we defined gene losses through gene CN < 2.0. The same thresholds were used to define gene losses in X chromosomes in female patients. Gene losses in X chromosomes in males were defined through gene CN < 1.0 for non-WGD samples and through gene CN < 1.5 for WGD samples. CNAs were unavailable for one relevant HMF sample and nine relevant TCGA samples, for which we assumed that no gene deletion events occurred.

### Pathway enrichment analysis

To understand the functional importance of the genes with SGMs of different mutational signatures, we performed an integrative pathway enrichment analysis using the ActivePathways method ^26^ (*FDR* < 0.05). The analysis was designed to prioritise genes and pathways that were enriched with signature-associated SGMs in multiple cancer types. We included the cancer types for which such genes were found, excluding colorectal cancer for which only one gene was found. For each cancer type, we selected the cohort with most cancer samples: lung (SBS4, TCGA), liver (SBS4, TCGA), breast (SBS13, HMF), head & neck (SBS13, TCGA), and uterine cancer (SBS13, TCGA). As the input to ActivePathways, we used a matrix of P-values of all protein-coding genes and the selected cancer types, such that each P-value reflected the enrichment of signature-associated SGMs in the gene and the cancer type. Gene sets of biological processes of Gene Ontology and molecular pathways of Reactome were derived from the GMT files in the g:Profiler web server ^55^ (downloaded January 3, 2022). Gene sets with 100-500 genes were analysed. Statistically significant pathways were selected (ActivePathways, FDR < 0.05). The results were visualised as an enrichment map ^56^ and the subnetworks were labelled interactively to find common functional themes of similar pathways and processes.

### Analysis of SGMs of SBS signatures in tumor subtypes and correlation with patient smoking history

We studied the number of signature-associated SGMs in each cancer genome in the representative cancer types (SBS4 in lung, SBS13 in breast; SBS18 in colorectal), and compared primary cancers in TCGA and metastatic cancers in HMF. Mean numbers of signature-associated SGMs per cancer genome were reported with 95% confidence intervals, by also including the samples where these SBS signatures were not detectable. We also compared the per-tumor SGM counts separately in various subtypes of lung and breast cancer. Subtype analysis was not performed in colorectal cancer due to limited subtype information available. Cancer subtype annotations for PCAWG were retrieved from the ICGC data portal, from patient information tables for HMF, and for TCGA from the TCGAbiolinks R package ^57^ (v. 2.18.0). Samples with unknown and missing subtype annotations were excluded. To validate the associations of SGMs and SBS signatures in the relevant cancer subtypes, we repeated the signature enrichment analysis of SGMs in samples of specific cancer subtypes using Fisher’s exact tests as described above. We also analysed SGMs of the tobacco signature SBS4 in the context of smoking history of lung cancer patients. We compared the subsets of TCGA cancer samples based on the four categories of patient smoking history that were derived from TCGAbiolinks. We compared two categories of SGMs: SGMs assigned to SBS4, and SGMs causing Glu > Stop substitutions. Non-parametric Wilcoxon rank-sum tests were used to compare mutation counts per patient in the four categories of smoking history. We performed one analysis by combining all lung cancer patients based on their smoking history, and two additional analyses focused on the two major histological subtypes (adenocarcinoma and squamous cell carcinoma). In breast cancer samples, we associated the frequency of SGMs per cancer genome with the gene expression levels of APOBEC enzymes *APOBEC3A* and *APOBEC3B*. We analysed breast cancer datasets in TCGA and HMF using matching RNA-seq datasets. Cancer samples with no SBS13 mutations were also included in the analyses. We excluded cancer samples with no matching RNA-seq data. Samples were split (median-dichotomised) into two subsets based on the median mRNA abundance of the APOBEC genes. The resulting two groups were compared using Poisson exact tests to compare mutation counts per cancer genome. Two types of mutations were considered: all snSNVs of the SBS13 signature, and all stop codon substitutions involving glutamic acids and serines combined (Glu > Stop, Ser > Stop).

## Acknowledgments

This work was supported by the Project Grant of Canadian Institutes of Health Research (CIHR) to J.R., the Operating Grant of the Cancer Research Society (CRS) to J.R., as well as the Investigator Award to J.R. from the Ontario Institute for Cancer Research (OICR). J.R. was supported by the New Investigator Award of the Terry Fox Research Institute (TFRI). A.T.B. was partially supported by an Ontario Graduate Scholarship. D.P. was partially supported by an Ontario STAGE HostSeq fellowship. Funding to OICR is provided by the Government of Ontario. This publication and the underlying study have been made possible partly based on the data that Hartwig Medical Foundation has made available to the study. The results published here are in part based on data generated by The Cancer Genome Atlas Research Network.

## Author contributions

N.A. led the data analysis, interpreted the results, and prepared the figures. K.C., M. S., A.T.B., D.P., K.A., and D.S. contributed to the data analysis, interpretation of results, and writing. N. A. and J.R. wrote the manuscript with input from all authors. J.R. conceived and oversaw the project. All authors reviewed and edited the manuscript and approved the final version.

