## Supplementary figures for "Mutational processes of tobacco smoking and APOBEC activity generate protein-truncating mutations in cancer genomes"

**Supplementary information**

Adler *et al*. (2023)

| **Consolidated cancer type** | **n total** | **PCAWG n** | **PCAWG ST** | **PCAWG ST n** | **HMF n** | **HMF ST** | **HMF ST n** | **TCGA n** | **TCGA subtype** | **TCGA ST n** |
|---|---|---|---|---|---|---|---|---|---|---|
| **Biliary** | **140** | 33 | Biliary-AdenoCA | 33 | 76 | Biliary | 76 | 31 | Cholangiocarcinoma | 31 |
| **Bone** | **512** | 94 | Bone-Cart<br>Bone-Epith<br>Bone-Leiomyo<br>Bone-Osteosarc | 9<br>11<br>34<br>40 | 209 | Bone/Soft tissue | 209 | 209 | Sarcoma | 209 |
| **Breast** | **1636** | 211 | Breast-AdenoCa<br>Breast-DCIS<br>Breast-LobularCa | 195<br>3<br>13 | 729 | Breast | 729 | 696 | Breast invasive carcinoma | 696 |
| **Colorectal** | **909** | 42 | ColoRect-AdenoCA | 42 | 522 | Colon/Rectum | 522 | 345 | Colon adenocarcinoma<br>Rectum adenocarcinoma | 258<br>87 |
| **Esophagus** | **408** | 95 | Eso-AdenoCa | 95 | 135 | Esophagus | 135 | 178 | Esophageal carcinoma | 178 |
| **Head and neck** | **611** | 54 | Head-SCC | 54 | 60 | Head and neck | 60 | 497 | Head and Neck squamous cell carcinoma | 497 |
| **Kidney** | **898** | 186 | Kidney-ChRCC<br>Kidney-RCC | 43<br>143 | 123 | Kidney | 123 | 589 | Kidney Chromophobe<br>Kidney renal clear cell carcinoma<br>Kidney renal papillary cell carcinoma | 51<br>317<br>221 |
| **Liver** | **716** | 313 | Liver-HCC | 313 | 57 | Liver | 57 | 346 | Liver hepatocellular carcinoma | 346 |
| **Lung** | **1454** | 78 | Lung-AdenoCA<br>Lung-SCC | 33<br>45 | 417 | Lung | 417 | 959 | Lung adenocarcinoma<br>Lung squamous cell carcinoma | 487<br>472 |
| **Lymphoid** | **258** | 196 | Lymph-BNHL<br>Lymph-CLL<br>Lymph-NOS | 104<br>90<br>2 | 25 | Lymphoid | 25 | 37 | Lymphoid Neoplasm Diffuse Large B-cell Lymphoma | 37 |
| **Nervous system** | **1085** | 273 | CNS-GBM<br>CNS-Medullo<br>CNS-Oligo<br>CNS-PiloAstro | 38<br>140<br>18<br>77 | 71 | Nervous system | 71 | 741 | Brain Lower Grade Glioma<br>Glioblastoma multiforme | 434<br>307 |
| **Neuro-endocrine** | **281** | 80 | Panc-Endocrine | 80 | 124 | Neuro-endocrine | 124 | 77 | Adrenocortical carcinoma<br>Pheochromocytoma and Paraganglioma | 63<br>14 |
| **Ovary** | **323** | 110 | Ovary-AdenoCA | 110 | 151 | Ovary | 151 | 62 | Ovarian serous cystadenocarcinoma | 62 |
| **Pancreas** | **465** | 231 | Panc-AdenoCA | 231 | 83 | Pancreas | 83 | 151 | Pancreatic adenocarcinoma | 151 |
| **Prostate** | **962** | 196 | Prost-AdenoCA | 196 | 366 | Prostate | 366 | 400 | Prostate adenocarcinoma | 400 |
| **Skin** | **688** | 65 | Skin-Melanoma | 65 | 226 | Skin | 226 | 397 | Skin Cutaneous Melanoma | 397 |
| **Stomach** | **518** | 62 | Stomach-AdenoCA | 62 | 41 | Stomach | 41 | 415 | Stomach adenocarcinoma | 415 |
| **Uterus** | **477** | 41 | Uterus-AdenoCA | 41 | 57 | Uterus | 57 | 379 | Uterine Corpus Endometrial Carcinoma | 379 |

**Supplementary Figure 1. Overview of cancer types used in the study.** Annotations of cancer subtypes (ST) in the three cohorts (TCGA, PCAWG, HMF) were consolidated to 18 types of cancer by anatomical sites.

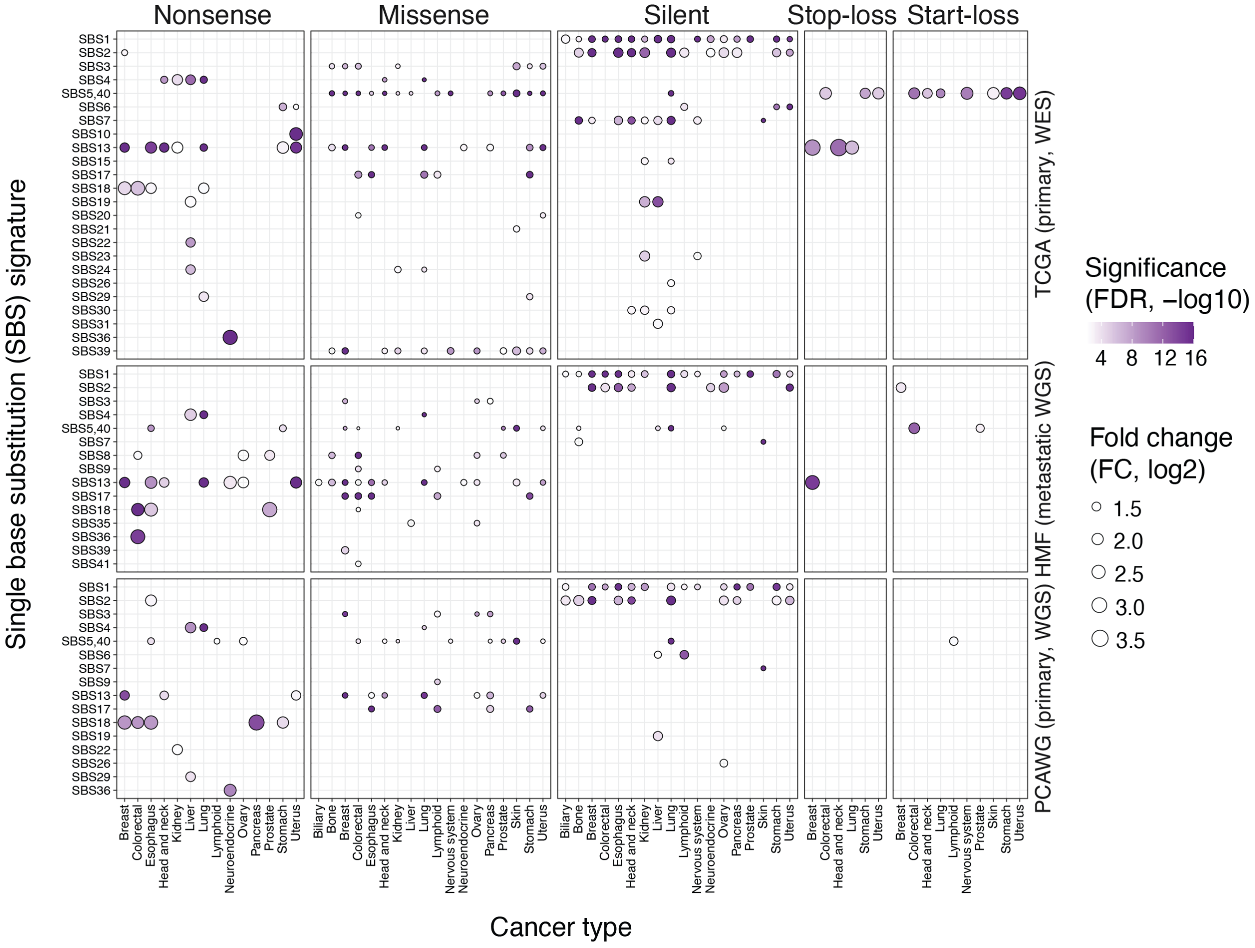


**Supplementary Figure 2. Protein-coding impact of mutational signatures in cancer genomes.** Landscape of mutational signatures that are enriched in functional classes of SNVs based on protein-coding impact. 18 types of primary and metastatic cancers were analysed separately. Significant associations are shown (Fisher's exact test; FDR < 0.01; FDR capped at $10^{-16}$). Fold-change (FC) shows the ratio of observed and expected SNV counts.

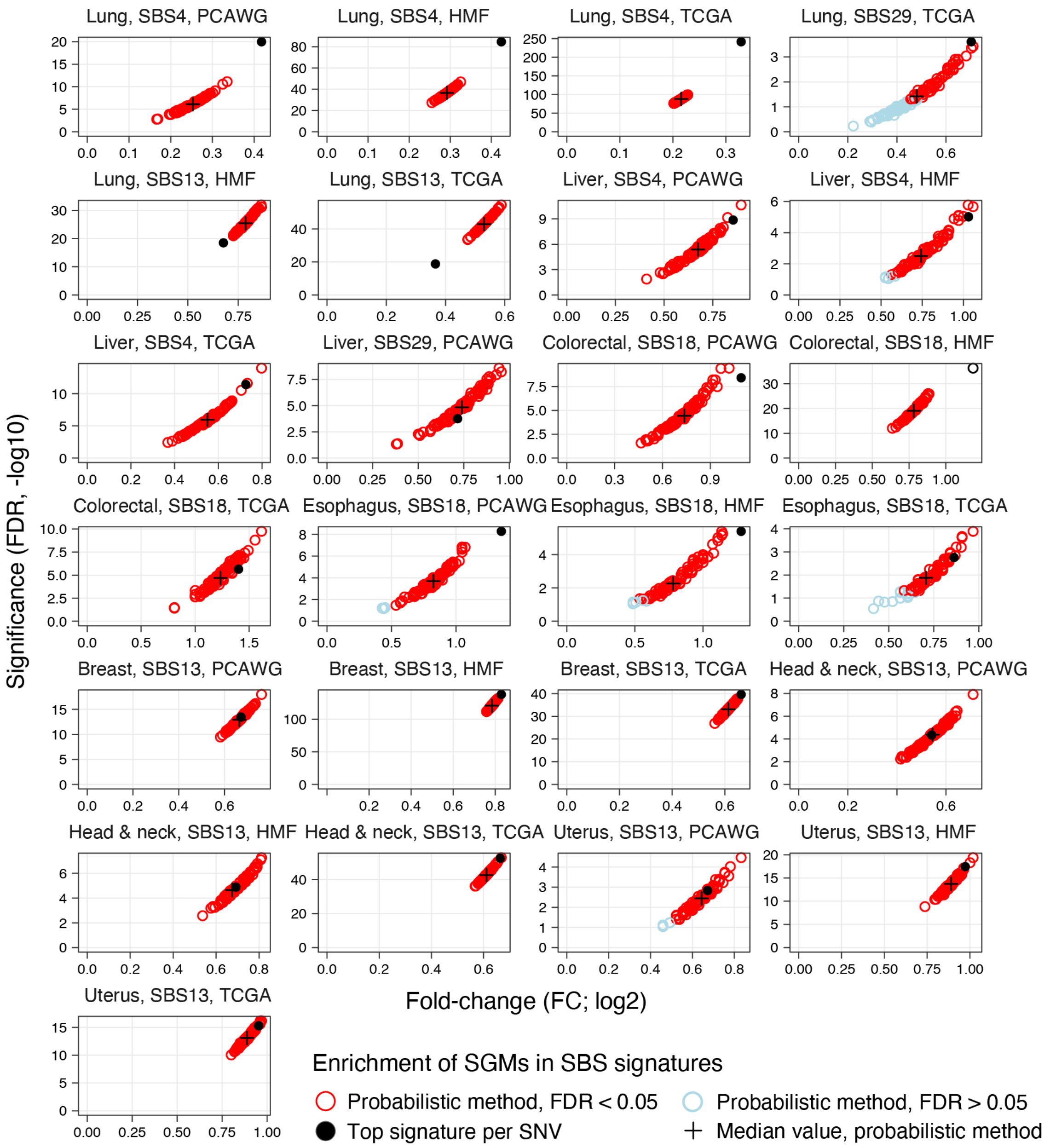


**Supplementary Figure 3. Probabilistic analysis of signature annotations of SNVs and enrichment analysis of SGMs.** Each SNV was assigned randomly to one SBS signature using the multinomial distribution parametrised by all SBS signatures in that cancer sample and the trinucleotide context of each SNV (*i.e.*, the probabilistic method). This procedure was repeated over 100 iterations. Each resulting set of SNVs was tested for enrichments of SGMs and the selected mutational signatures similarly to the analysis in Figure 1b-c. The probabilistic analysis of SBS signatures confirmed the main analysis of top-ranking signature annotations of SNVs as the enrichments of SGMs in SBS signatures remained highly significant.

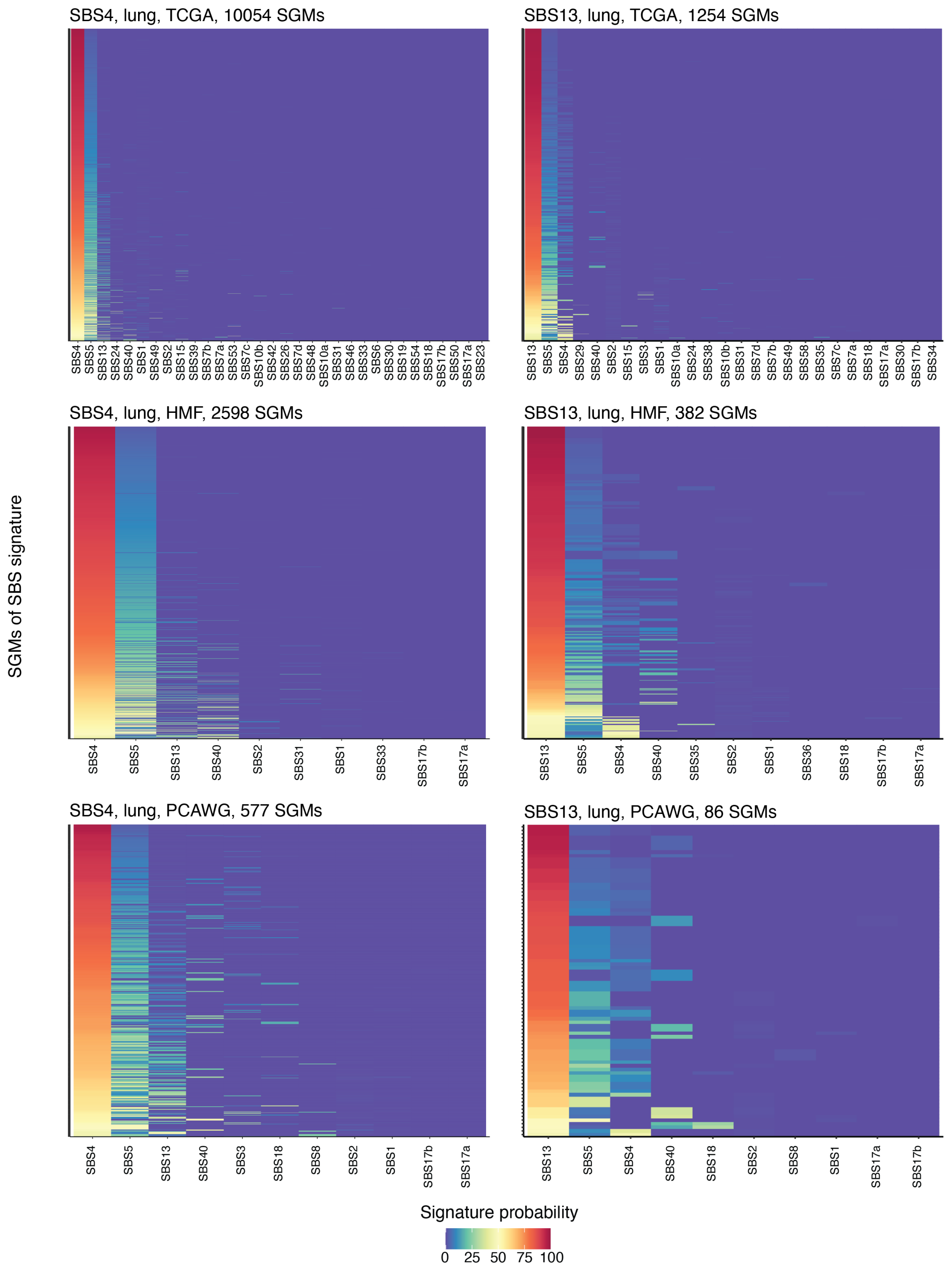


**Supplementary Figure 4. Comparison of signature annotation probabilities of SGMs of the signatures SBS4 and SBS13.** The columns are ordered by the cumulative probabilities of the SBS signatures. In lung cancer, the SNVs are often assigned to either the APOBEC signature SBS13 or the tobacco smoking signature SBS4 such that the two SBS signatures are the less-likely alternatives to each other, together with the clock-like signature SBS5 that has a relatively flat (featureless) profile.

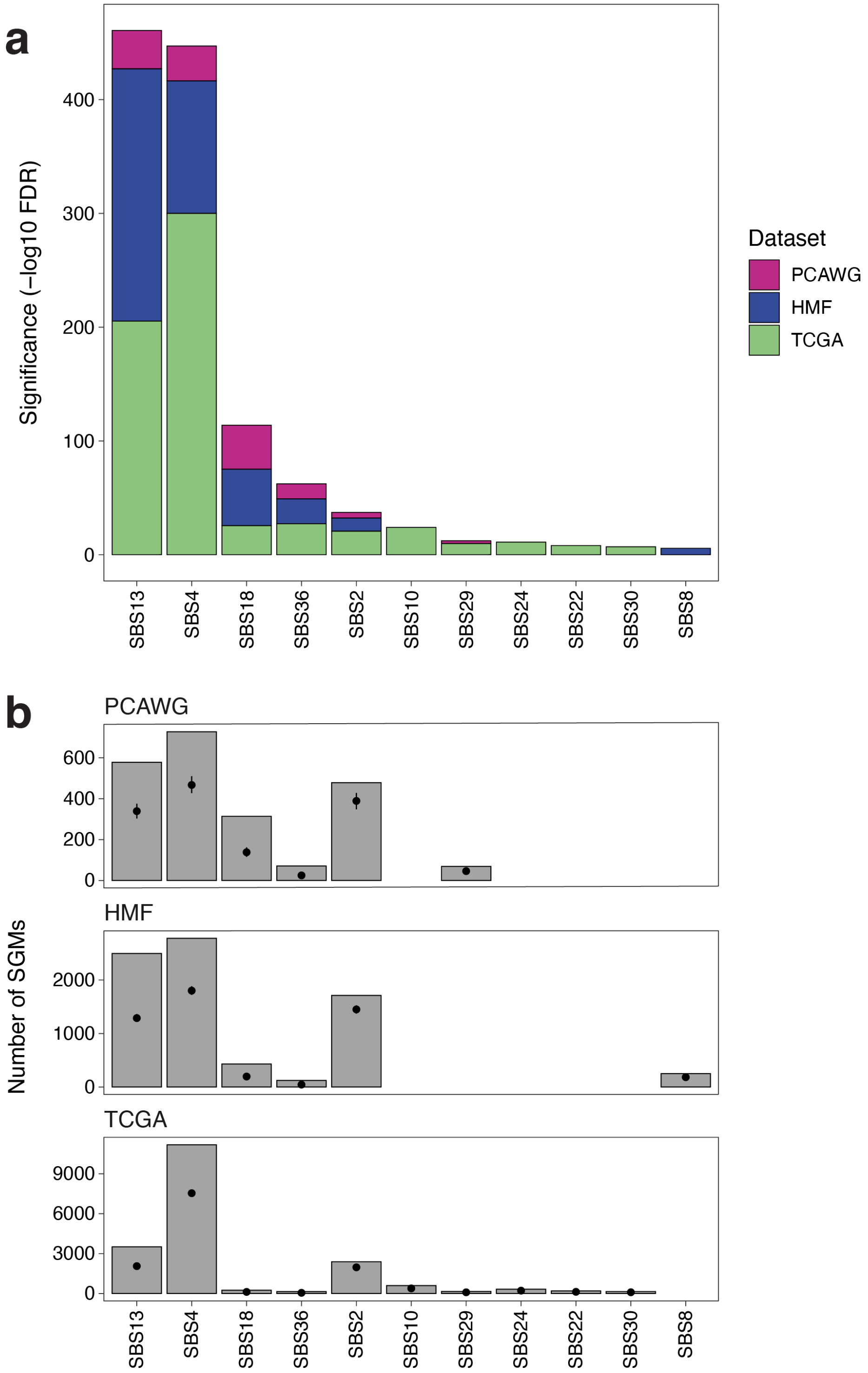


**Supplementary Figure 5. Pan-cancer enrichment of SGMs in mutational signatures. (a)** Cumulative significance of SGMs enriched in mutational signatures in the three cohorts (FDR-adjusted Fisher's exact tests, FDR < 0.01, capped $10^{-300}$). **(b)** Observed signature-associated SGMs in pan-cancer cohorts. Expected values were derived from binomial distributions and are shown with 95% Cis as points and whiskers.

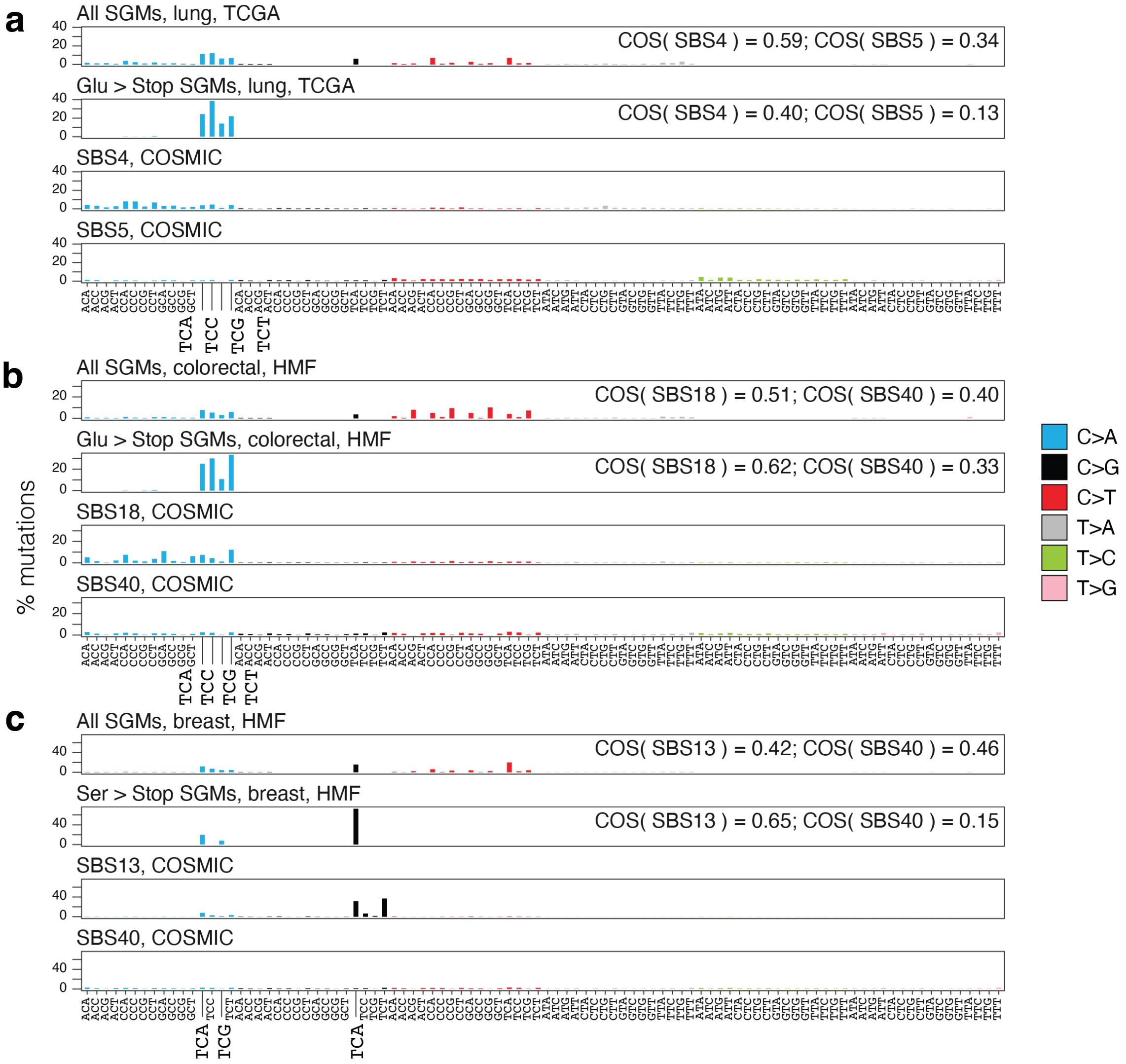


**Supplementary Figure 6. Analysis of COSMIC reference signatures and SGMs.** The bar plots show (i) observed SGMs, (ii) SGMs of the most abundant amino acid substitutions (Glu>Stop and Ser>Stop), (iii) reference COSMIC signatures enriched in SGMs (SBS4, SBS18, SBS13), (iv) and control signatures that represent the next most frequent signatures in the respective cancer types (clock-like signatures SBS5, SBS40). **(a)** The tobacco signature SBS4 in lung cancer. **(b)** The ROS signature SBS18 in colorectal cancer. **(c)** The APOBEC signature SBS13 in breast cancer. Cosine similarity (COS) scores of SGMs and reference signatures are shown at the top right.

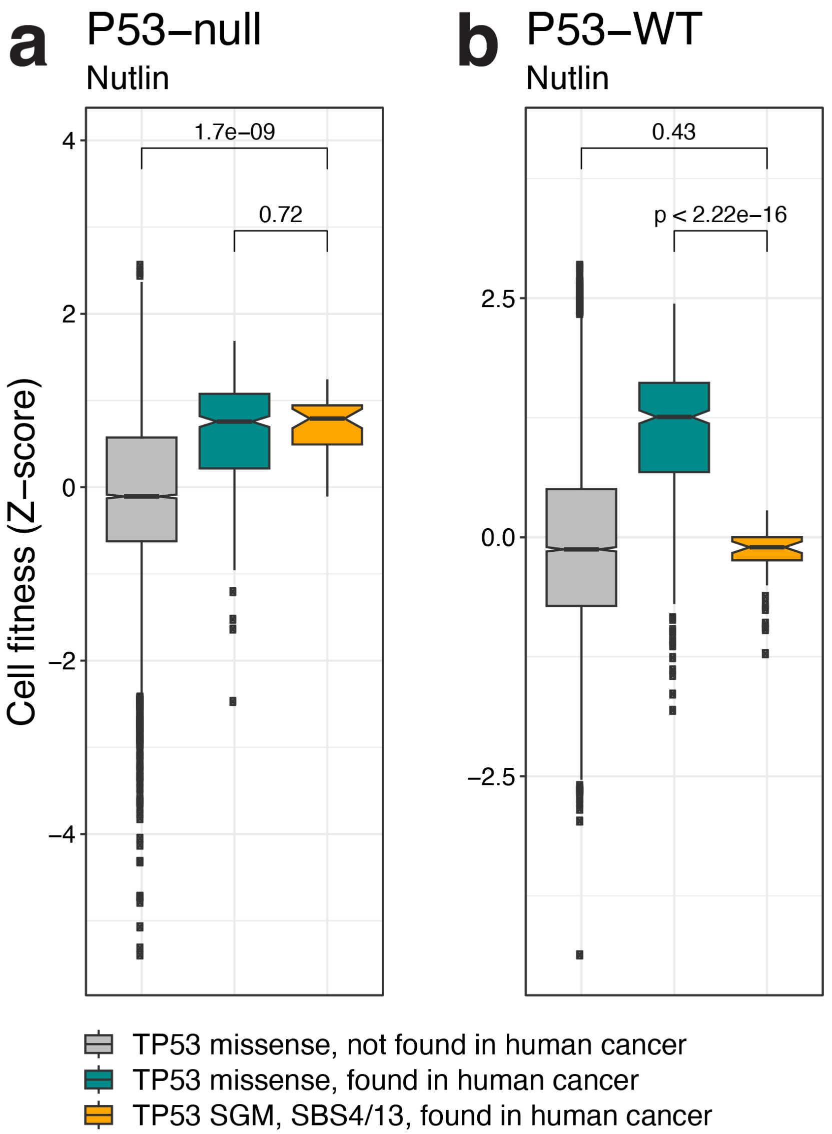


**Supplementary Figure 7. Functional analysis of SGMs in TP53.** SGMs of SBS4 and SBS13 signatures in TP53 in human cancer genomes are predicted to have loss-of-function effects on TP53, according to deep mutational scanning data from Giacomelli *et al*. (2018). **(a)** Cell fitness of TP53-NULL lung carcinoma epithelial cells (A549) transduced with SBS4/13 TP53 SGM mutants (yellow), TP53 missense SNV mutants found in cancer (teal) and all other possible missense SNV mutants (grey) *vs*. A549 cells transduced with wildtype TP53 upon nutlin treatment, which activates WT TP53 and results in a cell cycle arrest. Cells with SBS4/13 TP53 SGM mutant and TP53 missense SNV mutant exhibit higher fitness upon nutlin treatment indicative of p53 loss-of-function. **(b)** In TP53-wildtype cells, transduction of cancer-associated TP53 missense SNVs increases fitness, as most TP53 mutations found in cancer genomes are dominant negative and therefore inhibit the endogenous wildtype allele of TP53. Transduction of cancer-associated SGMs does not alter the fitness of p53-wildtype A549 cells, as these mutations represent bona fide LOF mutations without dominant negative functions. P-values from Wilcoxon rank-sum tests are shown.

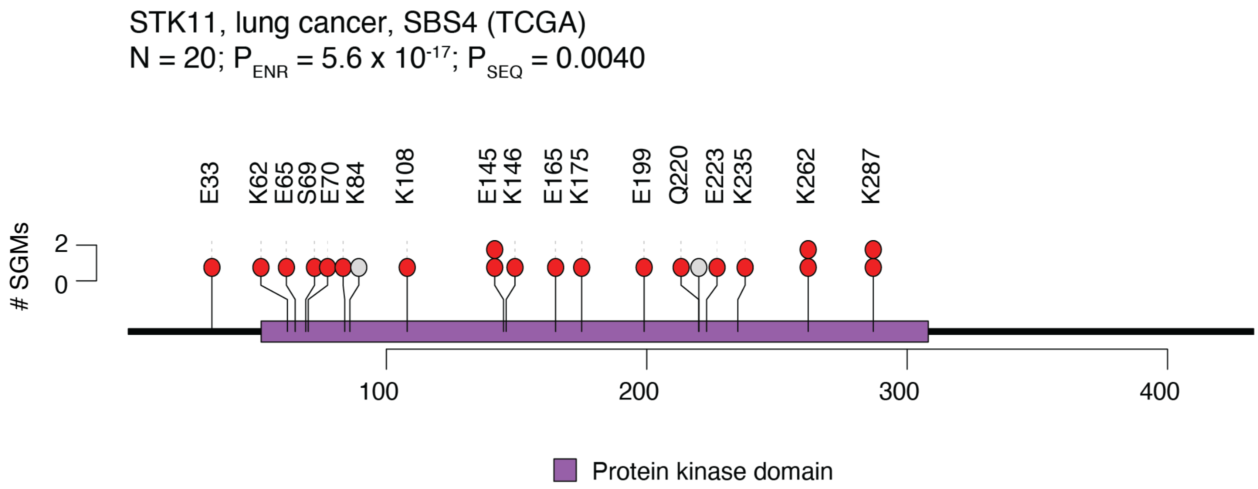


**Supplementary Figure 8.** SGMs in STK11 in lung cancer driven by the tobacco signature SBS4. Circles show all SGMs in the gene. SGMs are labelled with the reference residue and the protein sequence position, and colored red if they are derived from SBS4. The PFAM protein kinase domain is shown in purple. The title shows the number of SGMs associated with the signature ($N_{SBS}$), the P-value of signature-associated SGM enrichment ($P_{ENR}$, Fisher's exact test), and the over-representation of SGMs towards the start of the protein ($P_{SEQ}$, one-sample Wilcoxon rank-sum test).

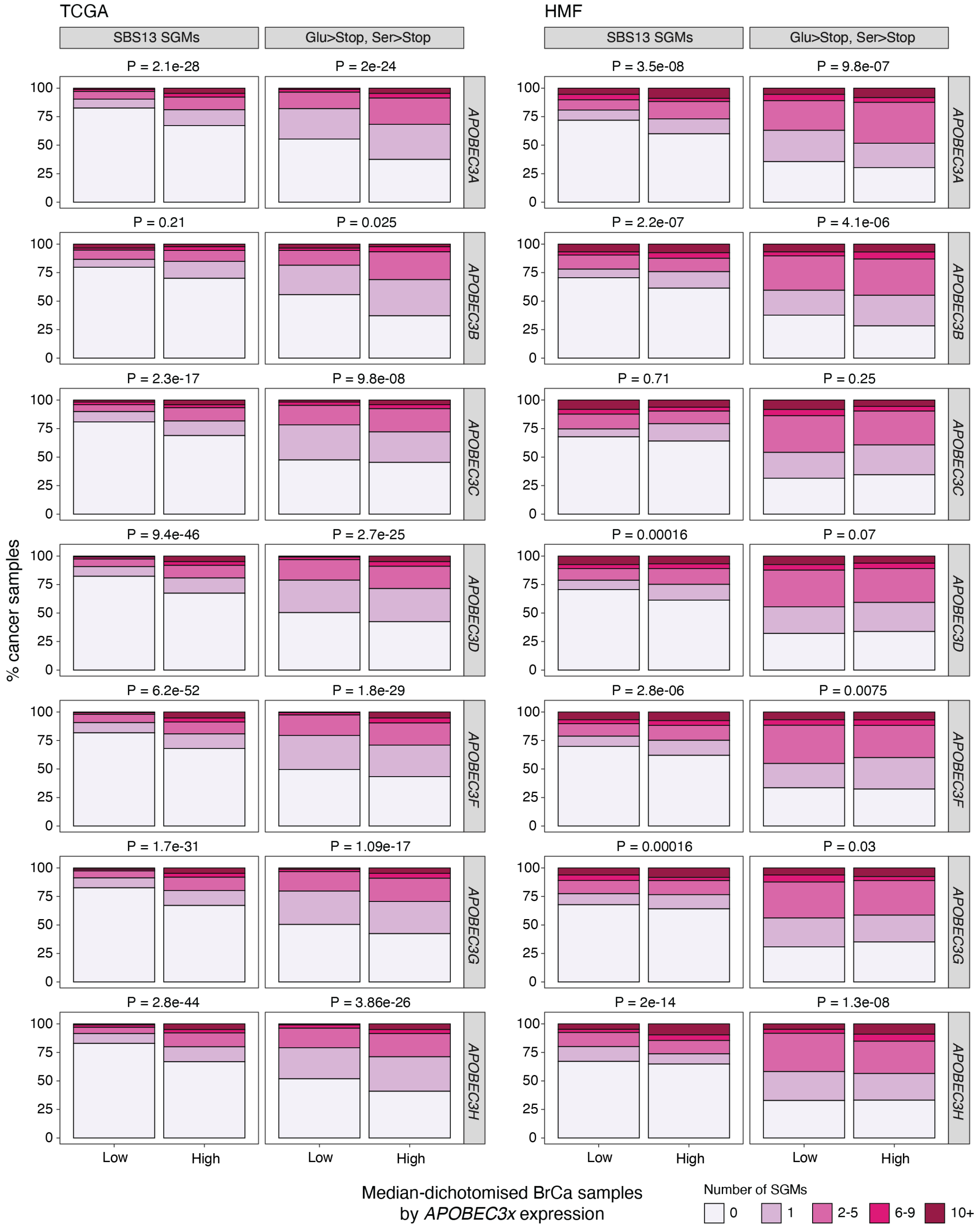


**Supplementary Figure 9.** SGM burden in breast cancer associates with increased expression of *APOBEC3* genes. Breast cancer samples were grouped into two equal sets based on median *APOBEC3* (*A-D,F-H*) expression and SBS13 SGMs and Ser>Stop and Glu>Stop substitutions were compared. P-values of Poisson exact tests are shown.
